# TreeFormer: A transformer-based tree rearrangement operation for phylogenetic reconstruction

**DOI:** 10.1101/2024.10.28.620561

**Authors:** Nhan Ly-Trong, Frederick A. Matsen, Bui Quang Minh

## Abstract

Phylogenetic inference is a fundamental problem in biology, which studies the origins and evolutionary relationships among species. Popular phylogenetic inference methods, such as IQ-TREE, RAxML, and PHYML, typically utilize heuristic tree search algorithms to seek a phylogenetic tree that maximizes the likelihood of the observed genetic data. However, tree search is time-consuming and often prone to local optima. To address these issues, we introduce TreeFormer, a new Transformer-based tree rearrangement operation for tree search. Experimental results show that TreeFormer achieves higher accuracy than FastTree 2 when reconstructing trees from real alignments with fewer than 1000 sites.

## 1 Introduction

### 1.1 Phylogenetics

Phylogenetic inference, which uses genetic data to reconstruct evolutionary trees, plays a vital role in understanding the evolutionary relationships among organisms (Felsenstein, 2004). Phylogenetic analysis is essential for exploring the Tree of Life, shedding light on the origins and evolutionary histories of species on the Earth (Misof et al., 2014; Jarvis et al., 2014; Weiss et al., 2016). Notably, phylogenetics is also crucial in health research, particularly in epidemiology. For instance, during the COVID-19 pandemic, phylogenetic tools were key to identifying the origins, tracking the spread, and monitoring emerging variants of the SARS-CoV-2 virus (Gorbalenya et al., 2020; Lu et al., 2020; Hodcroft et al., 2021; Vöhringer et al., 2021).

IQ-TREE (Minh et al., 2020) is a widely used maximum likelihood-based method for phylogenetic inference. Since its launch in 2013 (Nguyen et al., 2015), the core publications of IQ-TREE (Minh et al., 2013; Nguyen et al., 2015; Hoang et al., 2017; Kalyaanamoorthy et al., 2017; Minh et al., 2020) have accumulated over 45,000 citations on Google Scholar. Independent benchmarks have recognized IQ-TREE as the most accurate phylogenetic method (Zhou et al., 2018). IQ-TREE has significantly contributed to our understanding of SARS-CoV-2 transmission (Gao et al., 2020; Zhang et al., 2020), global threats like the Zika virus (Lessler et al., 2016), and the evolutionary relationships among animals (Simion et al., 2017), eukaryotes (Burki et al., 2016), and bacteria (Parks et al., 2018).

### 1.2 Machine Learning and Transformer

Machine learning can be defined as the task of approximating a function *y* = *f* (*x*), where *x* represents the input data and *y* denotes the expected output. If *y* consists of discrete categories, the task is referred to as classification, while if *y* takes continuous values, it is called regression (Duda & Hart, 1973). Unlike classical algorithmic methods which explicitly define the function *f* (*x*) through predefined rules, machine learning learns this relationship from large sets of training samples, called the training set. After training, the model is evaluated on unseen data, called the testing set, to assess its ability to generalize to new inputs (Mitchell, 1997).

A recent advancement in machine learning is the Transformer model, initially introduced by Vaswani et al. (2017) for natural language processing (NLP). Transformer is regarded as one of the most successful models in NLP, as highlighted by the success of ChatGPT (https://openai.com/chatgpt). In the context of NLP, the input data *x* is a sequence of words, where each word contributes to the overall meaning of the sequence at different levels; for example, content words like nouns and verbs are likely more important in conveying the message compared to function words like prepositions or determiners. The core of Transformer - attention mechanism, more specifically, self-attention, was designed to account for that. Given an input sequence *x* = {*s*_*i*_|*i* = 0, 1, 2, …, *L}*, of *L* words (or elements), the self-attention mechanism adjusts its focus (or attention) on these elements by using a triplet {*query, key, value}*. Specifically, self-attention mechanism computes the attention score *a*_*i*_ of the element *s*_*i*_ by considering all other elements *s*_*j*_ as

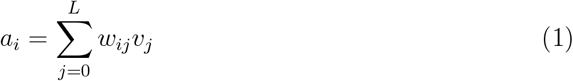

where:

- *w*_*ij*_ denotes the weight of information that is shared by an element *s*_*j*_ to *s*_*i*_.
- *v*_*j*_ is *value*(*s*_*j*_), which denotes the information of the element *s*_*j*_ to share with other elements. *w*_*ij*_ is computed from *query*(*s*_*i*_) and *key*(*s*_*j*_) as

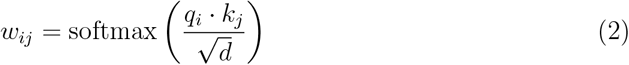

where
- softmax (Bridle, 1990) is a fundamental activation function in machine learning, commonly used to normalize a vector of *K* raw output values into a probability distribution across *K* output classes.
- *q*_*i*_ is *query*(*s*_*i*_).
- *k*_*j*_ is *key*(*s*_*j*_).
- *d* is the dimensionality of the query/key.

In recent years, the Transformer architecture has also been applied to biological sequence analysis (Jumper et al., 2021; Rives et al., 2021; Elnaggar et al., 2022).

### 1.3 Phyloformer

Phyloformer (Nesterenko et al., 2022) is a new transformer-based method for phylogenetic inference. Phyloformer takes a protein sequence alignment as input to predict the pairwise distances among the sequences as output, which is then fed into Neighbour Joining (NJ) (Saitou & Nei, 1987) to reconstruct a phylogenetic tree (Figure 1). Phyloformer comprises six attention layers (Nesterenko et al., 2022), allowing it to handle input alignments without constraints on the number of sequences or sites. However, due to resource and time limitations, Phyloformer was originally trained on simulated alignments with 20 sequences of 200 sites. Tested on simulated data, Phyloformer was about 25 times faster than the IQ-TREE software. With up to 20 sequences, Phyloformer was equally accurate as IQ-TREE (Nesterenko et al., 2022). However, its accuracy diminishes notably with larger alignments (*>* 20 sequences).

**Figure 1:**
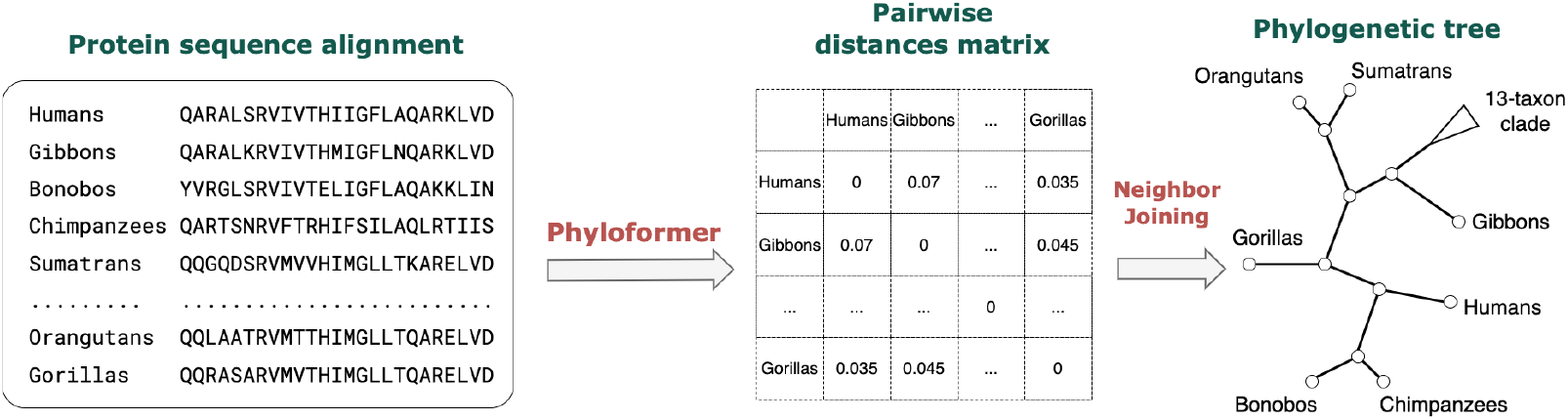
Illustration of reconstructing a phylogenetic tree from an MSA using Phyloformer and Neighbour Joining. Given an MSA with 20 protein sequences, Phyloformer predicts a 20×20 pairwise distance matrix, which serves as input for the Neighbour Joining to infer the phylogenetic tree. In the cartoon phylogenetic tree, 13 taxa were collapsed into a single clade for simplicity.

**Figure 2:**
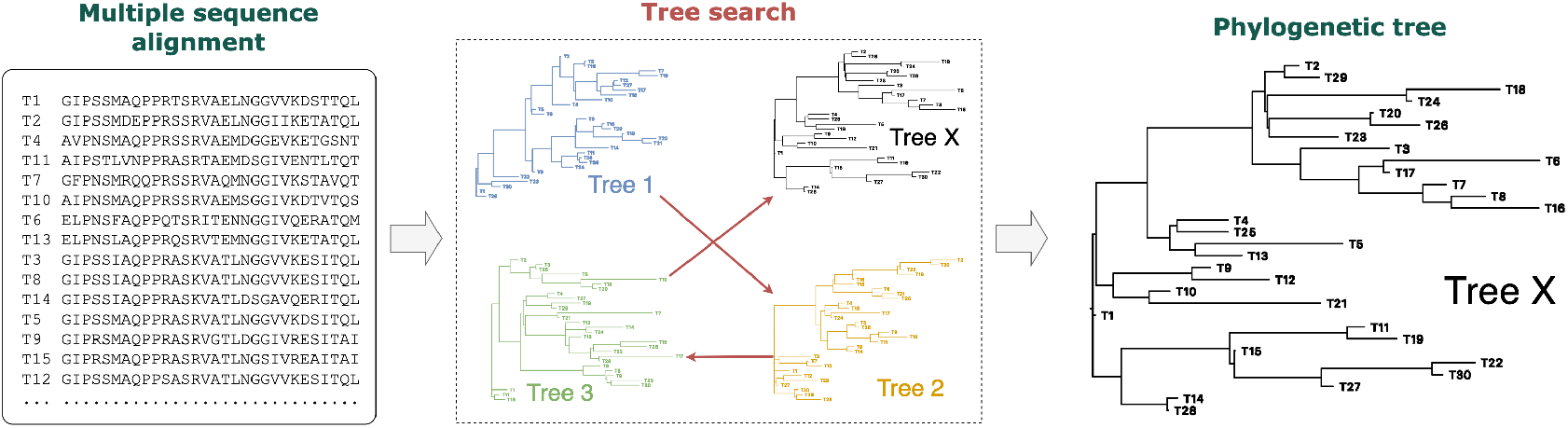
An example of inferring a phylogenetic tree from 30 sequences. Starting from an initial tree, Tree 1, we apply tree rearrangements to propose its neighboring trees. Among them, the tree with the highest likelihood, Tree 2, is accepted. This process is repeated with some heuristic strategies to overcome local optima. We then find Tree 3 before reaching the maximum likelihood tree, Tree X.

Maximum likelihood methods, such as IQ-TREE (Minh et al., 2020), RAxML (Kozlov et al., 2019), and PHYML (Guindon et al., 2010), commonly employ heuristic tree search algorithms to seek a phylogenetic tree that maximizes the likelihood of observing the input alignment. These heuristics explore the tree space by applying tree rearrangements such as Nearest Neighbour Interchange (NNI) or Subtree Pruning and Regrafting (SPR) (Felsenstein, 2004) that generate a set of neighboring trees from the current tree and evaluate the likelihoods of these trees. This process is repeated until no tree with a higher likelihood is found (Figure 2).

Tree search accounts for the majority amount of runtime and is susceptible to getting stuck in local optima. To mitigate these challenges, we propose TreeFormer, which extends Phyloformer to become a new machine learning-based tree rearrangement. Experimental results demonstrate that TreeFormer achieves higher accuracy than FastTree 2 (Price et al., 2010) when reconstructing trees from real alignments with fewer than 1000 sites. However, TreeFormer is still less accurate than IQ-TREE.

## 2 Methods

### 2.1 Overview of a new Transformer-based model (TreeFormer)

Figure 3 visualizes the idea of employing TreeFormer to propose a new tree, Tree T’, from the current tree, Tree T. We first randomly select a connected region formed by *S* subtrees or leaves. Then, we employ TreeFormer to predict the pairwise distances between subtrees and use Neighbour Joining to construct a new topology for that connected region. Finally, we re-connect the connected region to its subtrees to build Tree T’. During the tree search process, we can employ TreeFormer multiple times to optimize different connected regions of a large tree. In this research, we set *S* to 20 to ensure training feasibility, given the available computation and storage resources. While a larger connected region (i.e., a higher *S*) can reduce the runtime for optimizing a large tree, it requires significantly more computational resources and leads to an excessively longer training time. Besides, the original Phyloformer requires an MSA as input, which is inapplicable for a connected region because we cannot observe the sequences representing subtrees. To address that problem, we adopt partial likelihoods at the roots of these subtrees to compute the ancestral probabilities of each amino acid, which serve as input for TreeFormer.

**Figure 3:**
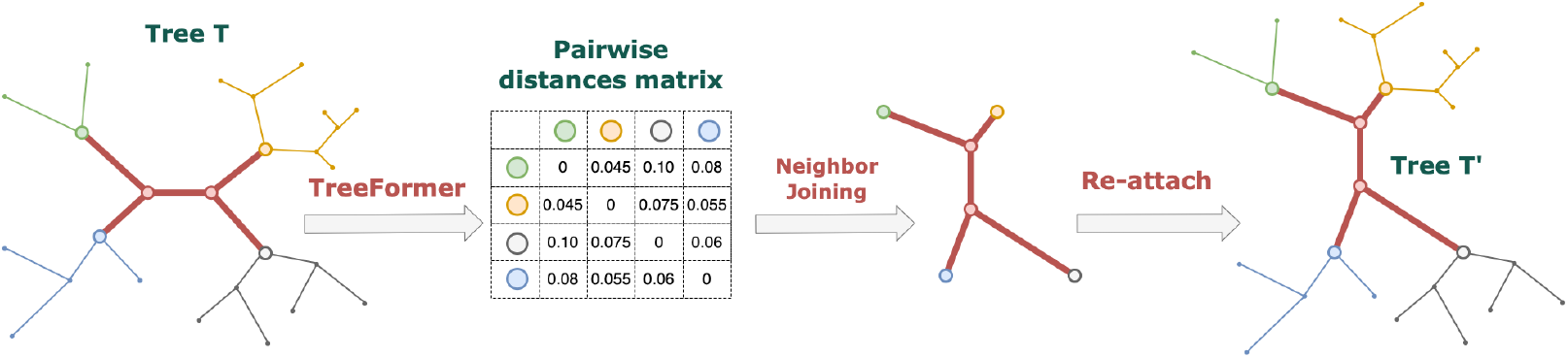
Illustration of using TreeFormer to propose Tree T’ from Tree T. We randomly select a connected region (which is formed by 20 subtrees or taxa). For simplicity, in this cartoon tree, the connected region is highlighted in red and is simplified to comprise only four subtrees colored in green, orange, black, and blue, respectively. The partial likelihoods at the roots of these subtrees are then fed into TreeFormer to predict the pairwise distances among those subtrees. Subsequently, a new topology for the connected region is constructed using Neighbor Joining. Finally, the connected region is reintegrated with its corresponding subtrees to obtain Tree T’.

In the following, we describe the four primary steps of the project: data generation, hyperparameter fine-tuning, model training, and performance evaluation.

### 2.2 Generating Data

Data plays a vital role in machine learning studies, which is critical to the success of the models. Therefore, in this project, we utilized both empirical and simulated data.

#### 2.2.1 Empirical Data

The EvoNAPS database (https://github.com/Cibiv/EvoNAPS) contains over 64,000 phylogenetic trees, and model parameters inferred by the IQ-TREE 2 software (v2.2.0.5) from about 22,600 DNA and 6,600 protein alignments. These alignments were gathered from three datasets: the PANDIT (Whelan et al., 2006), the OrthoMaM (Douzery et al., 2014), and the Lanfear (https://github.com/roblanf/BenchmarkAlignments/) databases.

In IQ-TREE, we developed new functions to randomly select a connected region comprising 20 subtrees from a given phylogenetic tree. For each subtree, the partial likelihood is computed and normalized so that the sum of the ancestral probabilities of amino acids at each site equals one. Gaps in the alignment are encoded as vectors with 20 equal elements (i.e., each element is 0.05).

From the EvoNAPS database, we extracted maximum-likelihood trees, which were reconstructed from amino acid alignments of at least 20 taxa. After removing short alignments (i.e., the number of sites is fewer than ten times the number of taxa) and abnormal trees with extremely long branches (*>* 10 units), we obtained 2,703 trees. For each tree with *n* taxa, we generated (*n* − 20)*λ* + 1 connected regions (data samples). *λ* is set at 1.5, so that we obtained 114,086 samples, which were then divided into a training set ( 90%) and a testing set ( 10%).

Similar to Phyloformer, to reduce the data size and the training time, we randomly subsampled (without replacement) 200 sites of the partial likehoods for all 102,601 samples in the training set. This means for each sample, 200 sites were randomly selected without preserving their original order. For the testing set, we subsampled 200 and 2000 sites, resulting in two testing sets: 200-site testing set (with 11,485 samples) and 2K-site testing set (with 2,201 samples). These datasets are hereby referred to as the “real-data training set”, “real-data 200-site testing set”, and “real-data 2K-site testing set”, respectively.

Furthermore, to evaluate the performance of TreeFormer on larger connected regions, we applied the above procedure to generate testing samples for connected regions comprising 20, 30, 40, and 50 subtrees. Regarding sequence length, we generated data with 200 sites and 2K sites. Each region size includes 1000 testing samples, resulting in a total of 8000 testing samples (i.e., 4 region sizes *×* 1000 samples per size *×* 2 sequence lengths). This testing set is called the “real-data larger region testing set”.

#### 2.2.2 Simulated Data

To simulate sequence alignments, we first estimated the best-fit distributions for model parameters from the EvoNAPS database using the Kolmogorov-Smirnov test implemented by Naser-Khdour et al. (2021). All model parameters and simulation settings are detailed in Table 1. We employed AliSim to simulate 3,780 alignments, from which 185,220 connected regions (samples) were extracted.

**Table 1:**
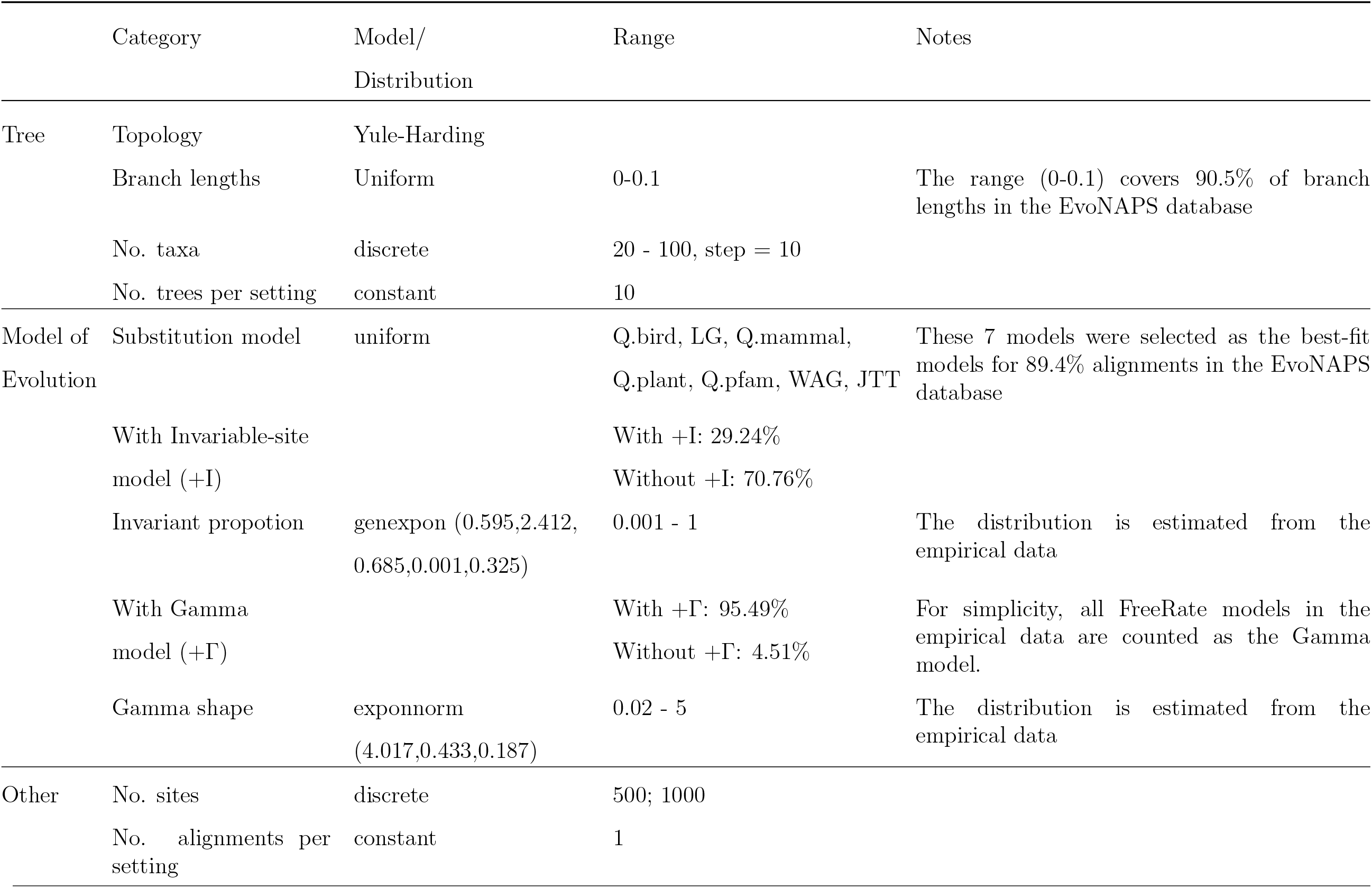
Model parameters and simulation settings for generating the simulated dataset.

All the partial likelihoods were then truncated to 200 sites by random subsampling (without replacement) of the sites. To reduce the computations for training, we further sampled 100,000 samples for the training set and 10,000 samples for the testing set. These datasets are referred to as the “simulated-data training set” and “simulated-data testing set”, respectively.

### 2.3 Fine-tuning Hyperparameters

Since Phyloformer was already well fine-tuned for its task, we hypothesized that the optimal settings for TreeFormer would be similar. Therefore, instead of fine-tuning the hyperparameters on a large parameter space, we tested a few revised settings of Phyloformer on a subset (25%) of the real-data training set. Results indicated that the best hyperparameter setting closely matches that of Phyloformer, as demonstrated in Table 2. The network architecture is identical to Phyloformer, with modifications only to the input layer (Figure 4).

**Table 2:** The Phyloformer’s, the tested, and the best hyperparameters.

| Hyperparameters | Phyloformers setting | Tested settings | Best setting |
| --- | --- | --- | --- |
| No. attention layers | 6 | 1, 3, 6, 9 | 6 |
| Dropout | 0 | 0, 0.1, 0.2, 0.3 | 0 |
| Batch size | 4 | 4, 8, 16, 32, 64 | 32 |
| amp (Automatic Mixed Precision) | Enable | Enable, Disable | Disable |
| Learning rate | 4e-4 | 4e-5, 4e-4, 4e-3 | 4e-4 |
| No. epochs | 185 | 185 | 185 |
| Early stop | Yes | Yes | Yes |
| Optimizer | Adam | Adam | Adam |

**Figure 4:**
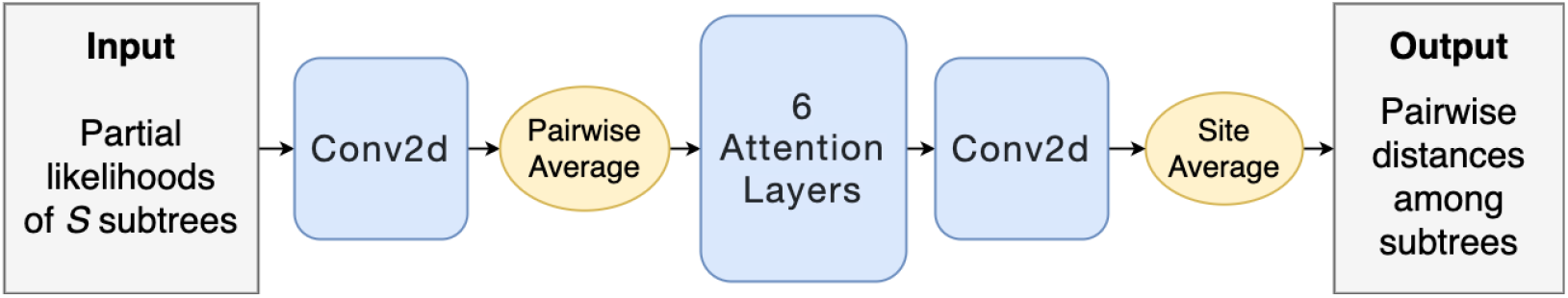
Network architecture of TreeFormer, the same as Phyloformer except the input layer. The input partial likelihoods of *S* subtrees are fed into a 2D convolution layer, and pairwise averages are applied over subtrees to convert the input into a three-dimensional tensor (*d × N × L*), where *d* is the size of the vector encoding the information for a site, set to 64 as per Phyloformer, 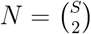 denotes the number of pairs of subtrees, and *L* is the sequence length. The tensor is passed through six attention blocks and a 2D convolution layer before applying an average over sites to compute a vector of *N* elements, representing the pairwise distances among subtrees.

### 2.4 Training Models

Given the best hyperparameter setting, we trained two machine learning models, TreeFormer RD and TreeFormer SD, independently using the real-data and the simulated-data training sets, respectively. We utilized the DataParallel library in Pytorch to parallelize the training. Each model took about 136 hours (wall time) to train using four Nvidia Tesla Volta V100-SXM2-32GB GPUs on the Gadi system (https://nci.org.au/our-systems/hpc-systems). Due to Gadi’s runtime limit, the training job was terminated and restarted from its checkpoint every 48 hours.

### 2.5 Evaluating the Models

#### 2.5.1 Evaluation metrics

To evaluate our models and benchmark TreeFormer against existing methods, we utilized the following five metrics.

1. The Pearson Correlation Coefficient between the predicted and true distances.
2. The Robinson Foulds (RF) distance (Robinson & Foulds, 1981) between the inferred and true connected regions.
3. The Pearson Correlation Coefficient between the inferred and true connected region lengths (i.e., the total branch length of each connected region).
4. The Branch Score Distance (BSD) (Kuhner & Felsenstein, 1994) between the inferred and true connected regions.
5. The runtimes of TreeFormer and existing methods, including Phyloformer, IQ-TREE 2 and FastTree 2.

The first three metrics were computed using Python libraries: *NumPy* for the Pearson Correlation Coefficients, and *ete3* for the RF distances and the connected region lengths.

The BSDs were calculated using the *treedist* tool (Felsenstein, 1989). The runtimes were measured using */usr/bin/time* in Linux.

#### 2.5.2 Evaluation scenarios

Our evaluation involves the following four testing scenarios. Their results are presented in the next section.

##### Testing TreeFormer RD and TreeFormer SD on the real and simulated testing subsets

To quickly compare TreeFormer RD and TreeFormer SD, we randomly subsampled (without replacement) two subsets of the simulated-data and the real-data 200-site testing sets, each with 1000 samples. Then, we tested the two models on these testing subsets. Among the two models, we named the more accurate one as TreeFormer, and conducted further testings in the following.

##### Testing TreeFormer on the real-data testing sets of 200 and 2K sites

To evaluate the effects of longer data on the performance of TreeFormer, we tested it on both the real-data 200-site and 2K-site testing sets.

##### Testing TreeFormer on real data with different sizes of connected regions

To evaluate the performance of TreeFormer on larger connected regions, we tested it on the “real-data larger region testing set”. This dataset was generated from connected regions comprising 20, 30, 40, and 50 subtrees. Recall that these connected regions were extracted from larger trees (of the EvoNAPS database). Thus, the number of subtrees is no greater than the number of taxa in the original trees. Here, we want to evaluate the accuracy of TreeFormer in reconstructing connected regions of different sizes.

##### Benchmarking TreeFormer against Phyloformer, IQ-TREE 2 and FastTree 2

TreeFormer serves as a novel tree rearrangement operation that can be invoked multiple times during the tree search process to optimize different connected regions of a large phylogenetic tree. However, integrating TreeFormer into existing software, such as IQ-TREE, requires substantial time and effort, which is beyond the scope of our proof-of-concept project. Currently, TreeFormer can be employed to infer a single connected region composed of 20 subtrees or taxa.

To evaluate the performance of TreeFormer, we benchmarked it against Phyloformer, IQ-TREE 2, and FastTree 2 on alignments of 20 taxa to ensure a fair comparison. First, we subsampled 1000 alignments of 20 taxa from the EvoNAPS database. We used IQ-TREE to infer the maximum likelihood trees, which served as the ground truth for comparisons. From these alignments, we created five datasets by subsampling 200, 500, 1000, 1500, and 2000 sites. Finally, we ran TreeFormer, Phyloformer, IQ-TREE 2, and FastTree 2 to reconstruct trees from these datasets. TreeFormer and Phyloformer were tested on one CPU (Intel Xeon Platinum 8268 (Cascade Lake) 2.9 GHz CPU) and then on one GPU (Nvidia Tesla Volta V100-SXM2-32GB) independently.

## Results

### 3.1 Testing TreeFormer RD and TreeFormer SD on real and simulated testing subsets

Figure 5 compares the true and predicted distances from TreeFormer RD and TreeFormer SD. TreeFormer SD (trained on simulated data) performs well on simulated data but poorly on real data, with Pearson Correlation Coefficients of 0.949 and 0.466, respectively (Figure 5A and B). While TreeFormer RD (trained on real data) can make accurate predictions on both simulated and real data, with Pearson Correlation Coefficients of 0.860 and 0.975, respectively (Figure 5C and D). These results suggest that our simulated data represents only a subset of the parameter space from the real data.

**Figure 5:**
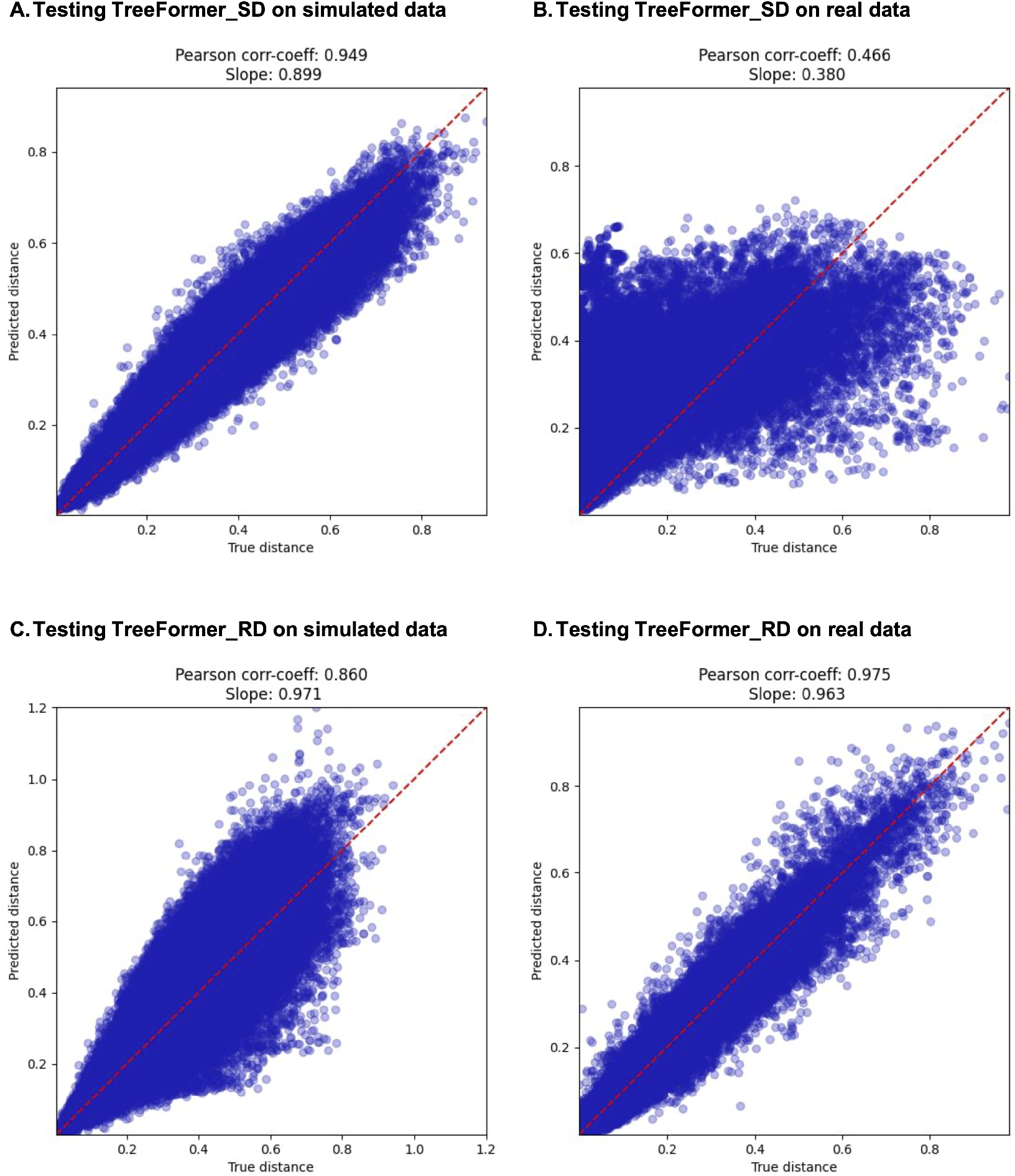
Comparison of the true and predicted distances of the two models: TreeFormer SD tested on simulated and real data (sub-panels A and B, respectively); and TreeFormer RD tested on simulated and real data (sub-panels C and D, respectively).

In the following, we refer to TreeFormer RD as TreeFormer and present additional results tested on this model.

### 3.2 Testing TreeFormer on the real-data testing sets of 200 and 2K sites

Figure 6 shows the testing results of TreeFormer on the real-data testing sets of 200 and 2K sites. For the 200-site dataset, the Pearson Correlation Coefficients between the true and predicted distances and that for the connected region lengths are 0.974 and 0.988, respectively (Figure 6A and B). The mean normalized RF distance and the mean BSD between the true and inferred connected regions are 0.539 and 0.068. For the 2K-site data, these coefficients increase to 0.992 and 0.995, respectively (Figure 6C and D), with mean normalized RF distance and mean BSD of 0.288 and 0.057.

**Figure 6:**
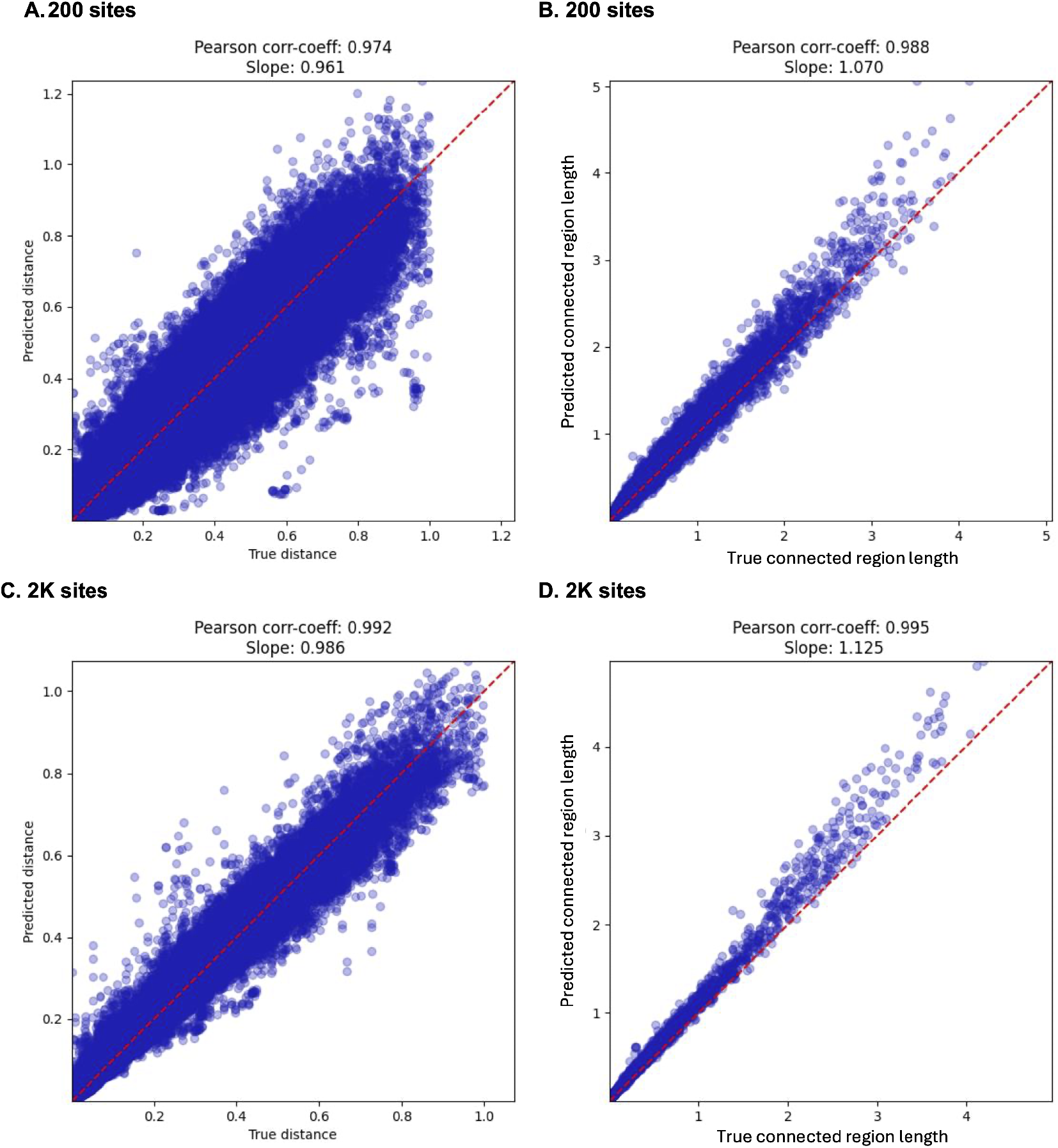
Testing results of TreeFormer on the real-data testing set of 200 (sub-panels A and B) and 2K sites (sub-panels C and D): The true and predicted distances (sub-panels A and C), and The true and inferred connected region lengths (sub-panel B and D).

In short, TreeFormer obtains higher accuracy on the 2K-site testing set than the 200-site set, which is expected as longer data provides more evolutionary signal.

### 3.3 Testing TreeFormer on real data with different sizes of connected regions

Figure 7 shows the mean normalized RF distances and the mean BSDs between the true and inferred connected regions by TreeFormer. As the size of the connected regions increases, the accuracy of TreeFormer decreases. Specifically, increasing the size of the connected regions from 20 to 50 subtrees results in the mean RF distance increasing from

**Figure 7:**
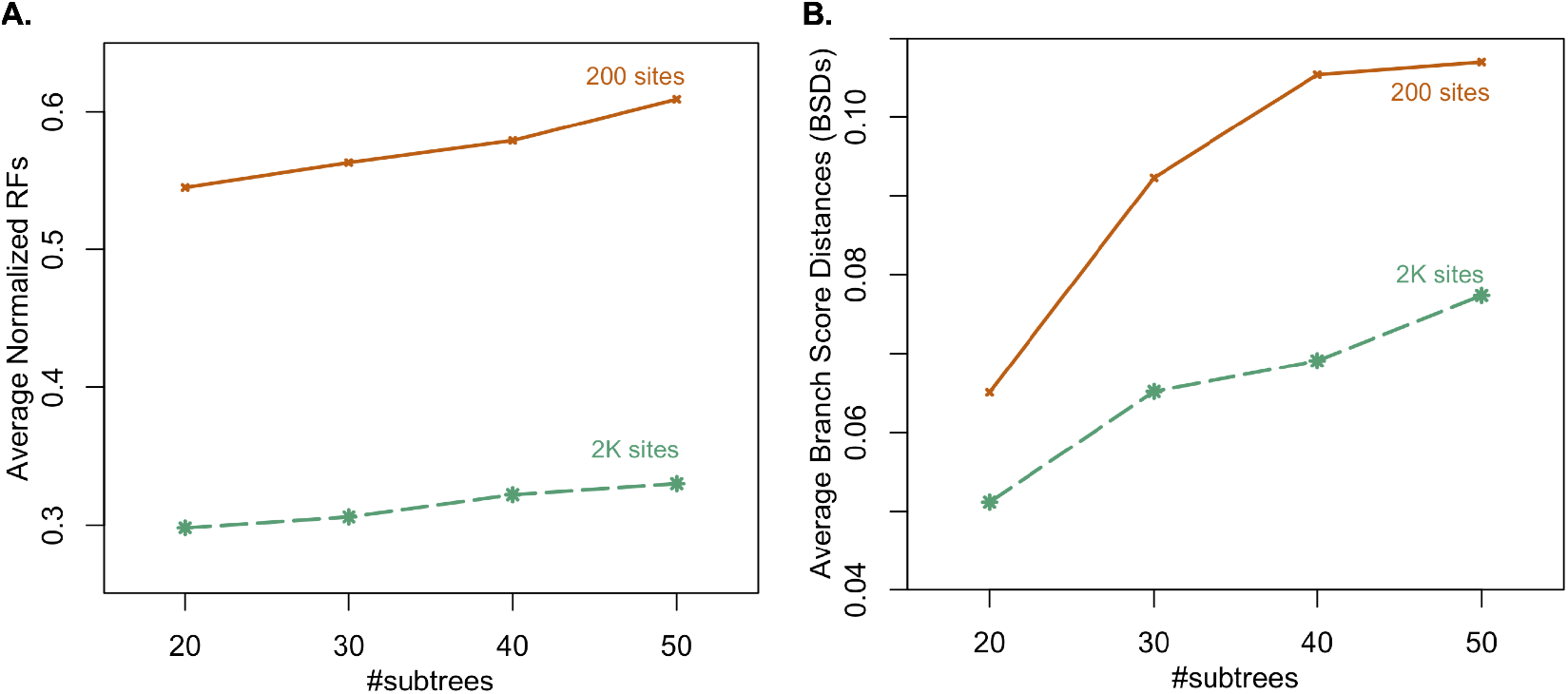
The mean RF distances (sub-panel A) and the mean BSDs (sub-panel B) between the truth and the inferred results of TreeFormer when testing on connected regions comprising 20, 30, 40, and 50 subtrees.

0.545 to 0.609 for 200-site testing samples and from 0.298 to 0.330 for 2K-site samples (Figure 7A). Similarly, the mean BSD increases from 0.065 to 0.107 for 200-site samples and from 0.051 to 0.077 for 2K-site samples (Figure 7B). These results suggest that TreeFormer can still yield reasonable results for larger connected regions (e.g., comprising 30, 40, or 50 subtrees). However, to obtain the most accurate result when reconstructing a large tree, we should invoke TreeFormer multiple times to optimize different connected regions formed by 20 subtrees, the size on which TreeFormer was trained. Additionally, these results corroborate our previous findings: longer data (e.g., 2K sites) significantly improves the accuracy of TreeFormer.

### 3.4 Benchmarking TreeFormer against Phyloformer, IQ-TREE 2 and FastTree 2

Figure 8 illustrates the mean normalized RF distances and the mean BSDs between the true trees and those inferred by TreeFormer, Phyloformer, IQ-TREE 2, and FastTree 2 from 20-taxon alignments with 200, 500, 1000, 1500, and 2000 sites. In general, increasing the sequence length improves the accuracy of all methods. CPUs and GPUs yield almost identical results for each machine-learning model. Among all methods, IQ-TREE 2 is the most accurate, achieving mean normalized RF distance and BSD of 0.123 and 0.026, respectively, for alignments with 2000 sites. TreeFormer outperforms FastTree 2 on short alignments before FastTree 2 takes over when the sequence length approaches 1000 sites. For alignments of 2000 sites, TreeFormer achieves mean normalized RF distance and BSD of 0.303 and 0.072, respectively, while FastTree 2 records corresponding values of 0.212 and 0.065. Phyloformer performs poorly, with mean normalized RF distance and BSD of 0.841 and 1.321 for alignments of 2000 sites. Note that the results were conducted using the initial version of Phyloformer, which was trained on non-gapped simulated data. This training set differs significantly from the real data used in our tests. Recently, a newer version of Phyloformer, with several improvements, has been introduced (Nesterenko et al., 2024). We expect the updated version to perform better.

**Figure 8:**
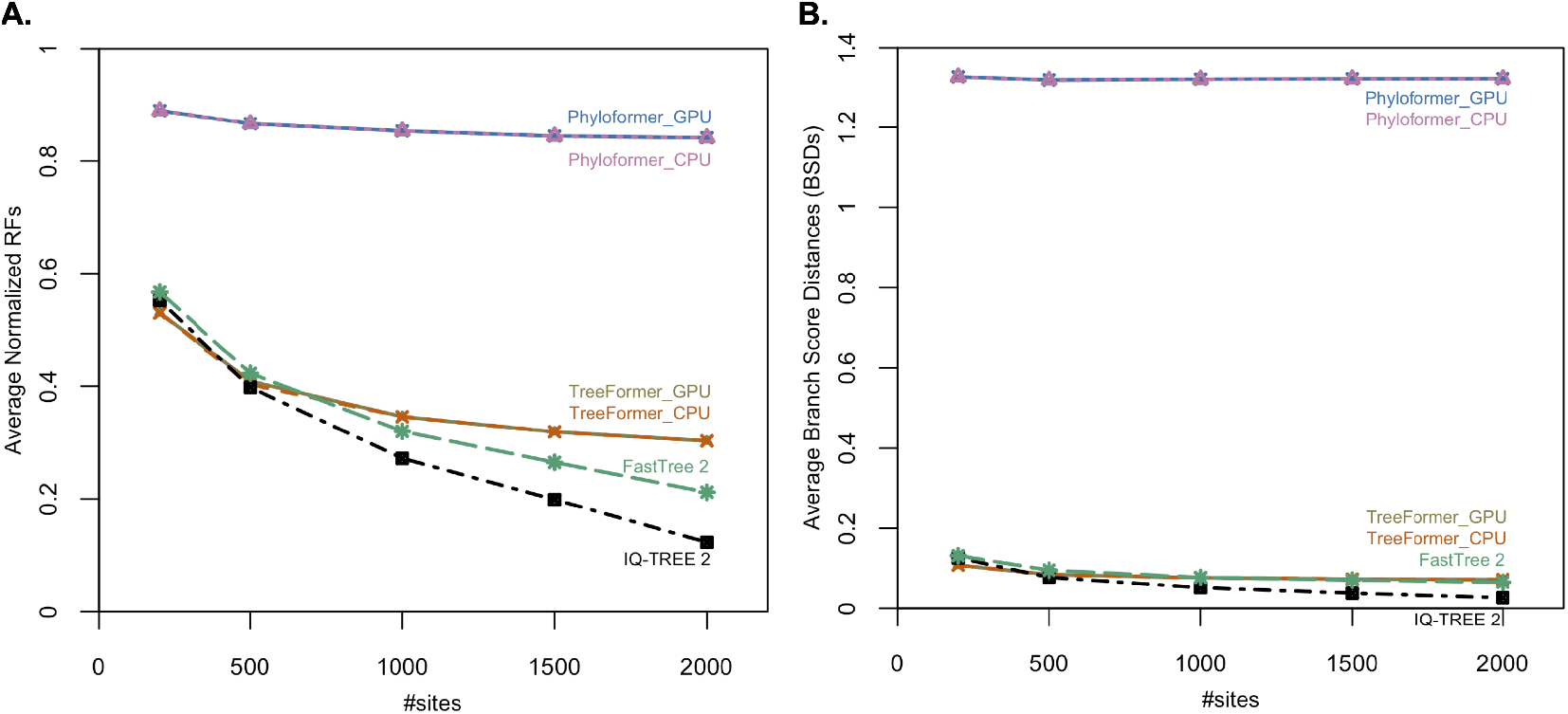
The mean RF distances (sub-panel A) and the mean BSDs (sub-panel B) between the truth and the trees inferred by TreeFormer, Phyloformer, IQ-TREE 2, and FastTree 2 from 20-taxon alignments of 200, 500, 1000, 1500, and 2000 sites. The results of TreeFormer and Phyloformer are reported for both cases tested on one CPU and on one GPU.

Figure 9 shows the runtimes (in minutes, log scale) of TreeFormer, Phyloformer, IQ-TREE 2, and FastTree 2. IQ-TREE required considerably more time than the other methods. For 1000 alignments of 2000 sites, IQ-TREE took a total of 1.6 days, while TreeFormer (using GPU), Phyloformer (using GPU), and FastTree 2 finished in 5, 6, and 28 minutes. When running on a CPU, TreeFormer and Phyloformer became slower, taking 6.3 and 6.7 hours, respectively. Note that we excluded the pre-processing time of executing IQ-TREE 2 to convert the alignments into an appropriate input format for TreeFormer, making TreeFormer slightly faster than Phyloformer. In our tests, that pre-processing step took 10 to 13 times longer than TreeFormer’s runtime. For instance, processing 1000 alignments of 2000 sites required 62 minutes in pre-processing alone - nearly 13 times longer than the runtime of TreeFormer (using GPU).

**Figure 9:**
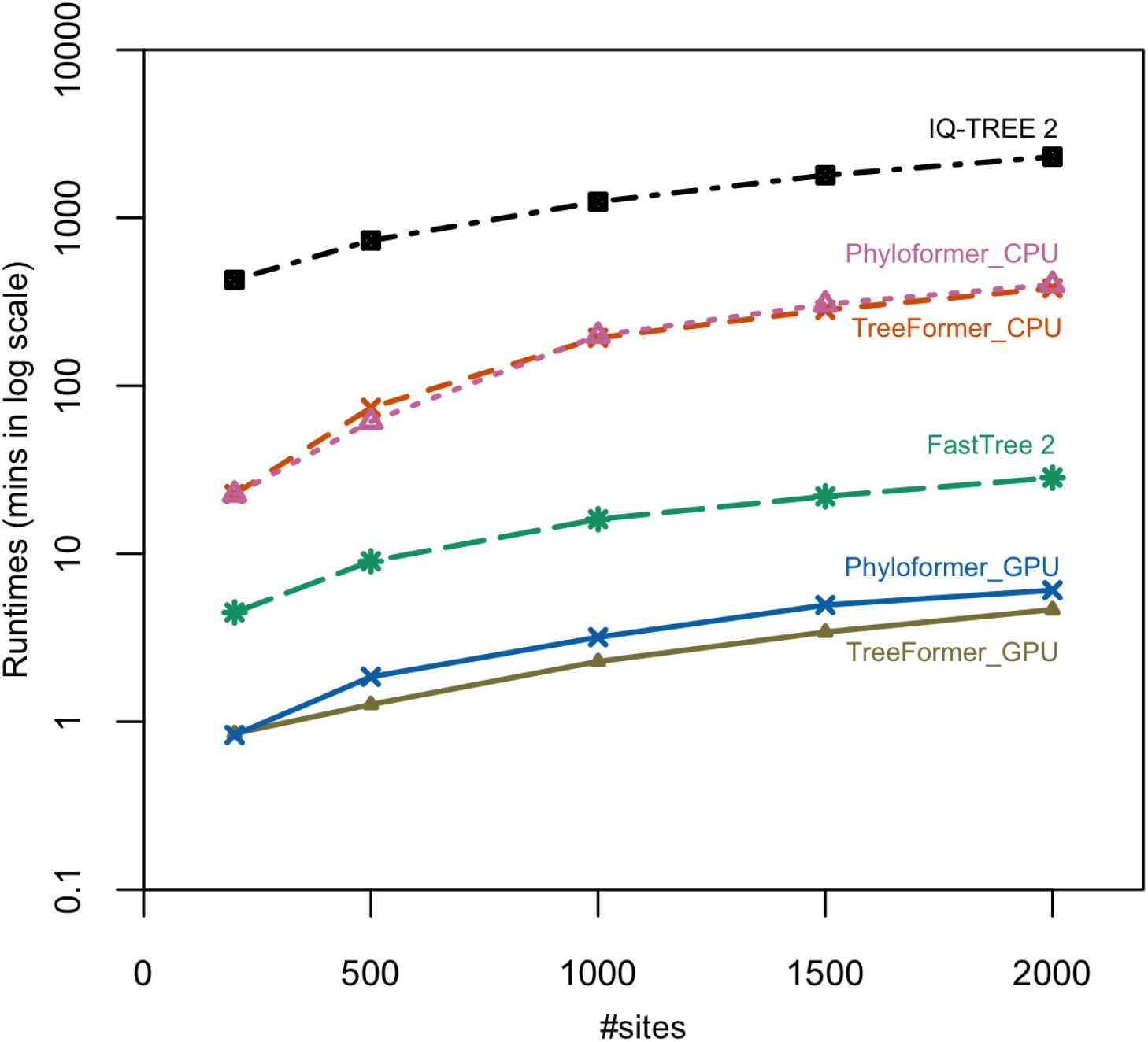
The runtimes of TreeFormer, Phyloformer, IQ-TREE 2, and FastTree 2 tested on 1000 alignments of 200, 500, 1000, 1500, and 2000 sites. The results of TreeFormer and Phyloformer are reported for both cases tested on one CPU and on one GPU.

## 4 Discussion

We present TreeFormer, which estimates the evolutionary distances among 20 subtrees from their partial likelihoods. TreeFormer can be employed iteratively or in parallel to optimize different subregions in a large tree, thus boosting the tree reconstruction. Experiments demonstrate that TreeFormer is more accurate than FastTree 2 in reconstructing trees from real data with fewer than 1000 sites. Additionally, increasing the number of sites of the input significantly enhances TreeFormer’s accuracy (figures 7 and 8).

However, due to limitations in research time and computational resources, TreeFormer is still in its infancy. Further improvements are necessary, including the following potential directions.

First, making the simulated data more realistic. We trained two models, TreeFormer SD and TreeFormer RD, on simulated and real data, respectively. Our results show that TreeFormer RD makes accurate predictions on both simulated and real data, while TreeFormer SD can only perform well on simulated data (Figure 5). The poor performance of TreeFormer SD on real data can be attributed to several reasons. Firstly, TreeFormer was designed to learn correlations between sequences and across sites. However, TreeFormer SD struggled to do so because sequence simulations assumed independent evolution across sites, thus preventing TreeFormer SD from capturing real-world dependencies. Secondly, our simulations did not account for complex biological processes such as gene duplication, gene transfer, recombination, indels, and natural selection pressures, which are likely presented in real data. Additionally, several key simulation parameters were too simplistic (Table 1), far different from the real data: tree topologies were generated from the Yule-Harding model, branch lengths followed a uniform distribution, and substitution models included only pre-estimated fixed-parameter models (e.g, Q.bird, LG, WAG, JTT, etc) without considering any flexibility, such as user-specified amino acid frequencies (e.g., Q.bird+F, LG+F, WAG+F, JTT+F, etc). To address this problem, we used real data for training in this research. If the amount of real data is limited, we can consider combining simulated and real data. To make the simulated data more realistic, we can use the special feature of AliSim to simulate new alignments that mimic real alignments (Ly-Trong et al., 2022).

Second, developing a hybrid tree search approach by combining machine learning (TreeFormer) and classical (NNI, SPR, or TBR) tree rearrangements together. To do so, we plan to integrate TreeFormer into an existing maximum likelihood method such as IQ-TREE. Both TreeFormer and NNI can be employed during the tree search of IQ-TREE, potentially avoiding local optima and accelerating the tree search process.

Third, improving the accuracy of TreeFormer by increasing the volume of training data and/or exploring alternative machine learning techniques. Transformer methods typically require a lot of training data. Due to the computational resource constraints, we trained our model with about 100,000 training samples. Retraining TreeFormer with a significantly larger dataset could further improve its accuracy. Another potential idea is employing ensemble learning which combines multiple machine learning models to improve the overall accuracy.

Fourth, testing TreeFormer on more empirical datasets. TreeFormer was trained and tested on real data extracted from the EvoNAPS database and may exhibit bias towards specific clades of species. Therefore, we also want to test TreeFormer on a more diverse range of data, such as bacterial and viral data, to assess the generalization of our model. Fifth, training TreeFormer for DNA data. While our current research focuses on protein data, extending TreeFormer or training a new model for DNA data would be beneficial. The EvoNAPS database includes a large number of DNA alignments, making this extension feasible with minimal revisions to the current codebase.

## Acknowledgements

This research was undertaken with the assistance of resources and services from the National Computational Infrastructure (NCI), which is supported by the Australian Government. The code was adopted from the Phyloformer repository (https://github.com/lucanest/Phyloformer). We thank Professor Robert Lanfear, Dr. Thomas Wong, Piyumal Demotte, and Hashara Kumarasinghe for their valuable suggestions and discussions.

